# Contrasting and Combining Transcriptome Complexity Captured by Short and Long RNA Sequencing Reads

**DOI:** 10.1101/2023.11.21.568046

**Authors:** Seong Woo Han, San Jewell, Andrei Thomas-Tikhonenko, Yoseph Barash

## Abstract

Mapping transcriptomic variations using either short or long reads RNA sequencing is a staple of genomic research. Long reads are able to capture entire isoforms and overcome repetitive regions, while short reads still provides improved coverage and error rates. Yet how to quantitatively compare the technologies, can we combine those, and what may be the benefit of such a combined view remain open questions. We tackle these questions by first creating a pipeline to assess matched long and short reads data using a variety of transcriptome statistics. We find that across datasets, algorithms and technologies, matched short reads data detects roughly 50% more splice junctions, with 10-30% of the splice junctions included at 20% or more are missed by long reads. In contrast, long reads detect many more intron retention events, pointing to the benefit of combining the technologies. We introduce MAJIQ-L, an extension of the MAJIQ software to enable a unified view of transcriptome variations from both technologies and demonstrate its benefits. Our software can be used to assess any future long reads technology or algorithm, and combine it with short reads data for improved transcriptome analysis.

## Introduction

Long reads sequencing technology has been revolutionizing genomic studies in recent years, leading to it being elected recently as “method of the year”^1^. The most commonly used platforms, Pacific Biosciences (PacBio) and Oxford Nanopore Technologies (ONT), both offer RNA sequencing with read length typically varying between a few hundred to a few thousand bases long, depending on the technology and the protocol used. Consequently, many algorithms have been developed for transcript discovery and quantification from long reads, such as FLAIR^2^, ESPRESSO^3^, IsoQuant^4^, Bambu^5^, TALON^6^, SQANTI^7^, and IDP^8^. Although both the technology and associated algorithms move at a fast pace, long reads RNA sequencing still suffers from several key limitations^9^. Specifically, many reads are still not long enough to capture entire transcripts, the high error rate makes it hard to detect exact isoforms and associated splice sites, and low coverage leads to limited isoform detection and quantification. In contrast, Illumina RNA short reads are typically only 100-150 bp long and therefore harder to assign to a specific isoform. Nonetheless, short reads still allow researchers to detect and quantify alternative splicing (AS) ‘events’ or, more generally, local splicing variations (LSV). LSV, first introduced in MAJIQ^10^, denote splits in a gene splice graph coming out of or into a reference exon. As such, LSV capture ‘classical’ AS events (e.g. cassette exons), but also more complex events involving multiple junctions or exons, including *de novo* (unannotated) junctions, exons, and introns. LSV are typically quantified using junction spanning reads in terms of percent spliced in (PSI, denoted Ψ), representing the relative percentage or ratio of isoforms with a specific splicing junction or intron retention (IR).

The availability of different short and long reads technologies raises the natural question of how these compare and whether can they be effectively combined. Yet previous work involving long reads has focused mainly on the benefits it may offer, lacking a comprehensive comparative evaluation of the resulting transcriptome maps. Similarly, tools that combine short and long reads to aid researchers in downstream splicing analysis are still underdeveloped.

To address these needs we developed an analysis pipeline and accompanying software, MAJIQ-L. The analysis pipeline shown in Figure 1-a takes as input three sources of information: Transcriptome annotation; short reads processed by MAJIQ V2^11^; and long reads in gtf format, processed by the user’s algorithm of choice. It then computes and displays an extensive set of statistics that contrast the available annotation and the two sequencing sources in terms of novel junctions, introns, coverage, inclusion levels, etc. such that existing gaps between the three sources can be captured (Figure 1-b). Using the three input sources, MAJIQ-L constructs unified gene splice graphs with all isoforms and all LSVs visible for analysis. This unified view is implemented in a new visualization package (VOILA v3), allowing users to inspect each gene of interest where the three sources agree or differ (Fig 1-c).

**Fig. 1:**
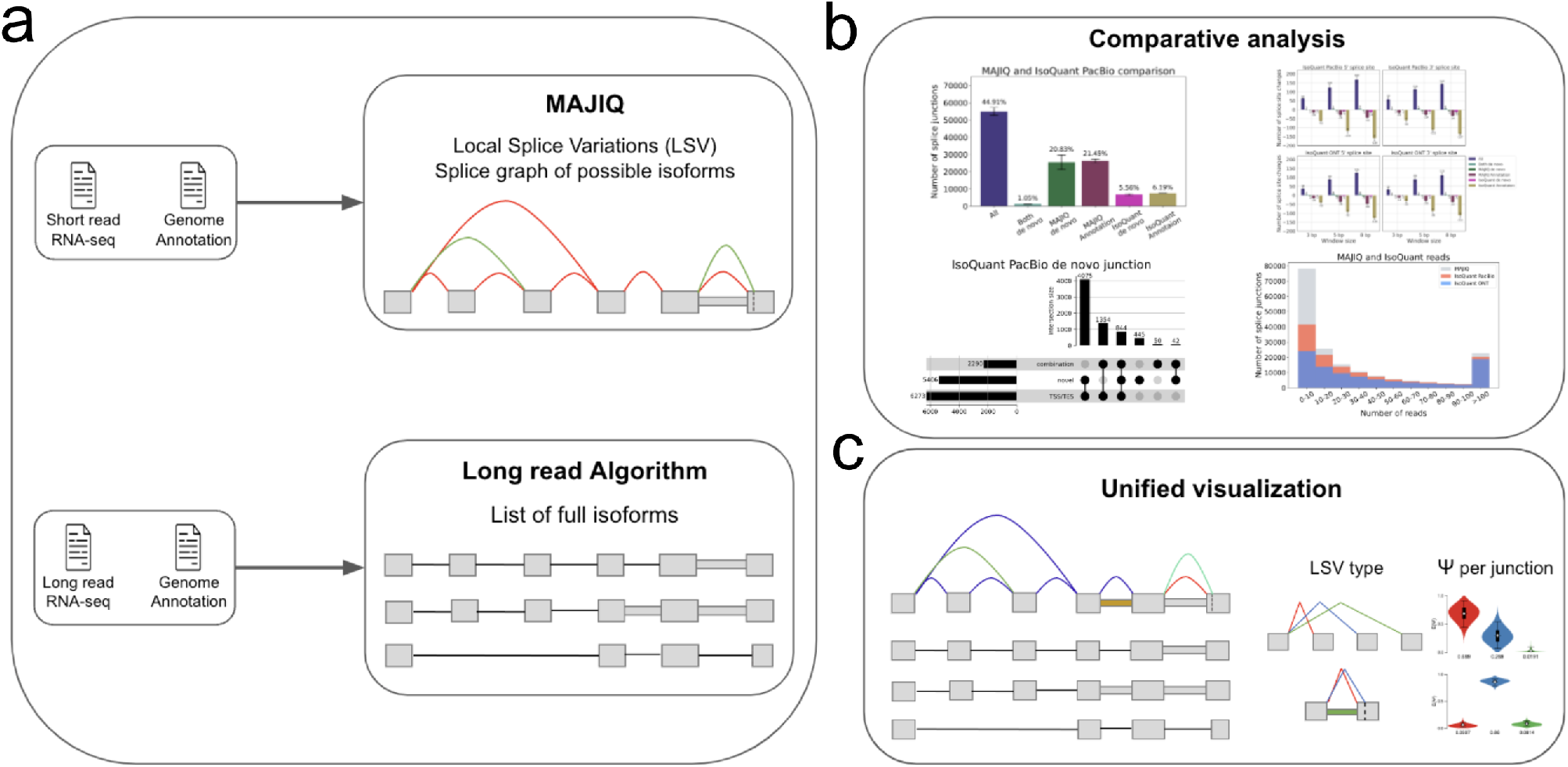
MAJIQ-L overview. **(A)** Input: Matched short and long reads RNA-sequencing data with genome annotation. Short reads are mapped with an aligner, then passed to MAJIQ, producing a splice graph with a matching set of LSV per gene. Long reads are processed with a long read algorithm (*e.g.*, IsoQuant) to produce isoforms counts. **(B)** The outputs from the two RNA sequencing sources along with the annotation are compared. Each splice junction or intron retention is assigned to each of the possible six categories depending on which subset of the three sources support it. Various statistics regarding the location, coverage, and overlap of those elements are computed to compare the three sources and explore the source of discrepancy (see main text). **(C)** MAJIQ-L includes a unified visualization package, VOILA v3, for downstream splicing analysis. It allows users to see which splice junction maps to which isoforms and where the different sources of information agree or disagree in detection and quantification.

We apply MAJIQ-L to matched short and long reads from several datasets involving both PacBio and ONT using four different long reads transcriptome mapping algorithms. First, we contrast short and long reads by statistics reflecting splice junction detection and quantification. Next, we inspect the coverage difference between short and long reads and 3’ to 5’ bias in long read sequencing. Finally, we turn to intron retention, showing that as expected long reads detect many more introns than short reads, but short reads detect longer introns. Finally, we demonstrate the usefulness of a combined long and short reads analysis using VOILA v3 for splicing variations in the SRSF11 gene.

## Materials and Methods

### Processing coverage differences between short and long reads

As the total number of bases sequenced across short and long reads technologies are different, we used Seqtk (https://github.com/lh3/seqtk) by sub-sampling from either short or long reads datasets before providing them as inputs to MAJIQ and long read tools. This step allows all platforms to have similar coverage for a fair comparative analysis. The coverage summary of pre and post-sub-sampled number of bases for each dataset can be found in Supplementary Table 1-3.

### MAJIQ’s short reads splicing analysis

We used STAR (v2.7.10b)^12^ to align short RNA-seq reads, performing a two-step gapped alignment to GRCh38, GENCODE release 42. MAJIQ then combines the annotation and aligned reads to build a splicegraph for each gene which includes *de novo* elements such as junctions, intron retention, and exons. The resulting splicegraphs are used as MAJIQ-L’s input along with long read tools’ output in gtf file format for the downstream comparative analysis.

### GTF file output files from long read tools

Long RNA-seq reads were mapped to GRCh38, GENCODE release 42, using minimap2 (v2.24)^13^ in splice mode. All long read tools were provided with the same BAM file, reference genome, and reference annotation. IsoQuant was run with the default parameters with the appropriate data type using ‘*−data type*’ option. ESPRESSO and FLAIR were launched with the default parameters in 30 threads. As Bambu outputs all reference transcripts, including unexpressed ones, we filtered out all transcripts with read count values that are smaller than 1 as the authors recommended. Software versions and command line options are in Supplementary Table 4.

### Inferring posterior distribution in long reads PSI per junction

The posterior distribution over PSI Ψ*_j_* for a junction *j* is modeled as a binomial distribution, where *r_j_*denotes long reads aligned to each junction *j* in the LSV:

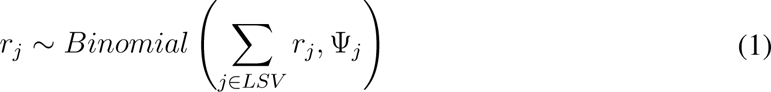

As in MAJIQ’s short reads model, we set a prior distribution on PSI which favors either high or low PSI values, which can be generalized by the Jeffrey’s prior for an LSV with *j* junctions:

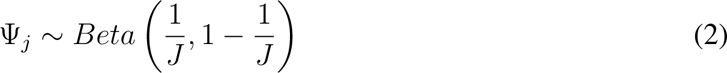

Since this prior is conjugate to the binomial distribution, our posterior distribution of Ψ*_j_* given the observed number of reads are the following:

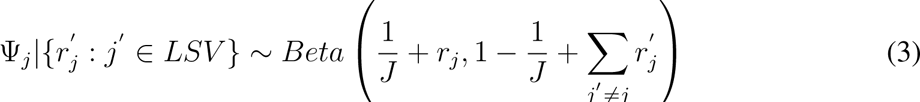

The resulting distributions over PSI for both short and long reads are shown as violin plots by the VOILA visualization package.

## Results

### Short reads detect 40-60% more splice junctions at the same coverage level

To contrast the observed transcriptome complexity by short and long reads, we compared splice junctions detected by short reads processed by STAR^12^ followed by MAJIQ’s LSV analysis^10^ and four different long read algorithms^2;3;4;5^ (see Methods for details). Each splice junction was assigned to one of six possible categories, depending on which source supported it (Fig 2a). This analysis was performed using three different datasets: Three replicates of human cell lines from the Long-read RNA-seq Genome Annotation Assessment Project (LRGASP) Consortium^14^ (shown in Fig 2b); Three heart atrial appendage samples from GTEx v9^15^ and a PDX cell line sample derived from a patient with a relapsed B Cell Acute Lymphoblastic Leukemia (B-ALL, Supplementary Fig 1). To address coverage difference, we sub-sampled the files such that the total number of bases sequenced across the various platforms was similar (see Materials and Methods). Overall, our results indicate that across all 3 datasets short reads detected 40-60% more splice junctions, with PacBio detecting about 10% more junctions than ONT. Some differences between long read algorithms were also apparent. FLAIR detected the most amount of long reads only splice junctions (*≈* 10%), while Bambu reported the least (*≈* 1.5%). These differences may reflect lower precision and recall respectively^4^. We also performed a comparative analysis when using the original dataset without sub-sampling, where long reads have 1.3-2.9 fold more bases sequenced than short reads (LRGASP, B-ALL) and where Illumina has 1.7-fold more coverage than ONT(GTEx). Similar trends were observed in this analysis as well (see Supplementary Fig 2, and Supplementary Table 1-3 for details).

**Fig. 2:**
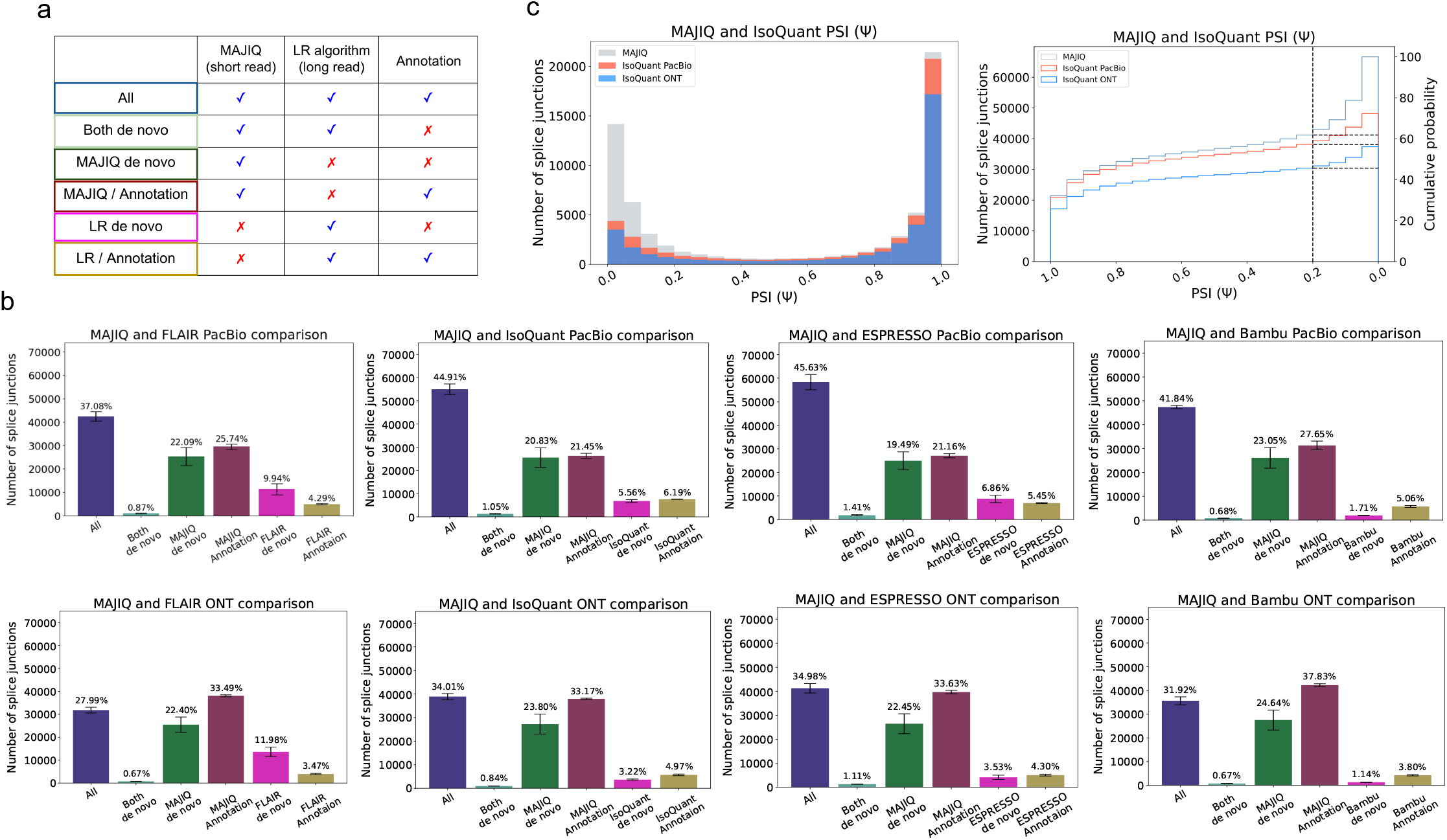
Splice junctions comparative analysis. **(A)** Any detected splice junction can fall into one of six categories, each represented by a color, depending which of the three sources of information (short reads, long reads, annotation) support it. **(B)** Bar charts corresponding to the aforementioned six categories. Mean and standard error bars are computed using matched datasets from three replicates of human cell-line sequenced by the LRGASP^14^. This data includes short reads processed by STAR and MAJIQ, Long reads from PacBio and ONT assays, and four long read algorithms used to process the long reads data. **(C)** Taking the splice junctions reported in (B) by MAJIQ (green) and assessing the number of those also identified when using PacBio (tomato) or ONT (blue) long reads, as a function of the PSI values. Here IsoQuant was used for long reads data. Note that if a junction appears in multiple LSV, the lowest PSI values are chosen (x-axis). The graph on the right is the CDF for the histogram shown on the left. Dashed lines denote splice junctions with a PSI of 20% or more.

The significant difference in splice junction detection naturally raises the question if the junctions uniquely detected by either technologies are real. In the case of short reads, MAJIQ only reports splice junctions when multiple split reads map to multiple positions, which has been shown to lead to very few false positives^16;17^. In contrast, long reads only splice junctions are more likely to involve false positives given their known high error rates and limited coverage. Regardless, the significant number of short reads only splice junctions begs the question of whether those additional junctions are meaningful. The results shown in Figure 2c for IsoQuant indicate most short reads only splice junctions are, as expected, relatively lowly included (Ψ *<* 10%). Nonetheless, the cumulative distribution function plot shows IsoQuant+PacBio misses 10% of junctions with significant inclusion levels (Ψ *>* 20%) and IsoQuant+ONT misses 30% of junctions detected by MAJIQ (Fig 2c right). When comparing the different algorithms, IsoQuant misses the least amount of junctions with Ψ *>* 20% and Bambu misses the most with the same inclusion level (Supplementary Fig 8b).

### Patterns of novel splice variants difference in short and long reads

Next we turned to better characterize the splice junctions that are unique to either short or long reads technologies. For short reads, we classified *de novo* junctions reported by MAJIQ into two categories: A junction involving novel splice sites, and a junction creating a novel combination of known splice sites (Fig 3a, top). Compared with IsoQuant, the distribution of the two categories is the same for PacBio and ONT, showing about 90% involve novel splice sites while the rest is novel combination. A similar trend is observed when compared with different algorithms (Supplementary Fig 3).

**Fig. 3:**
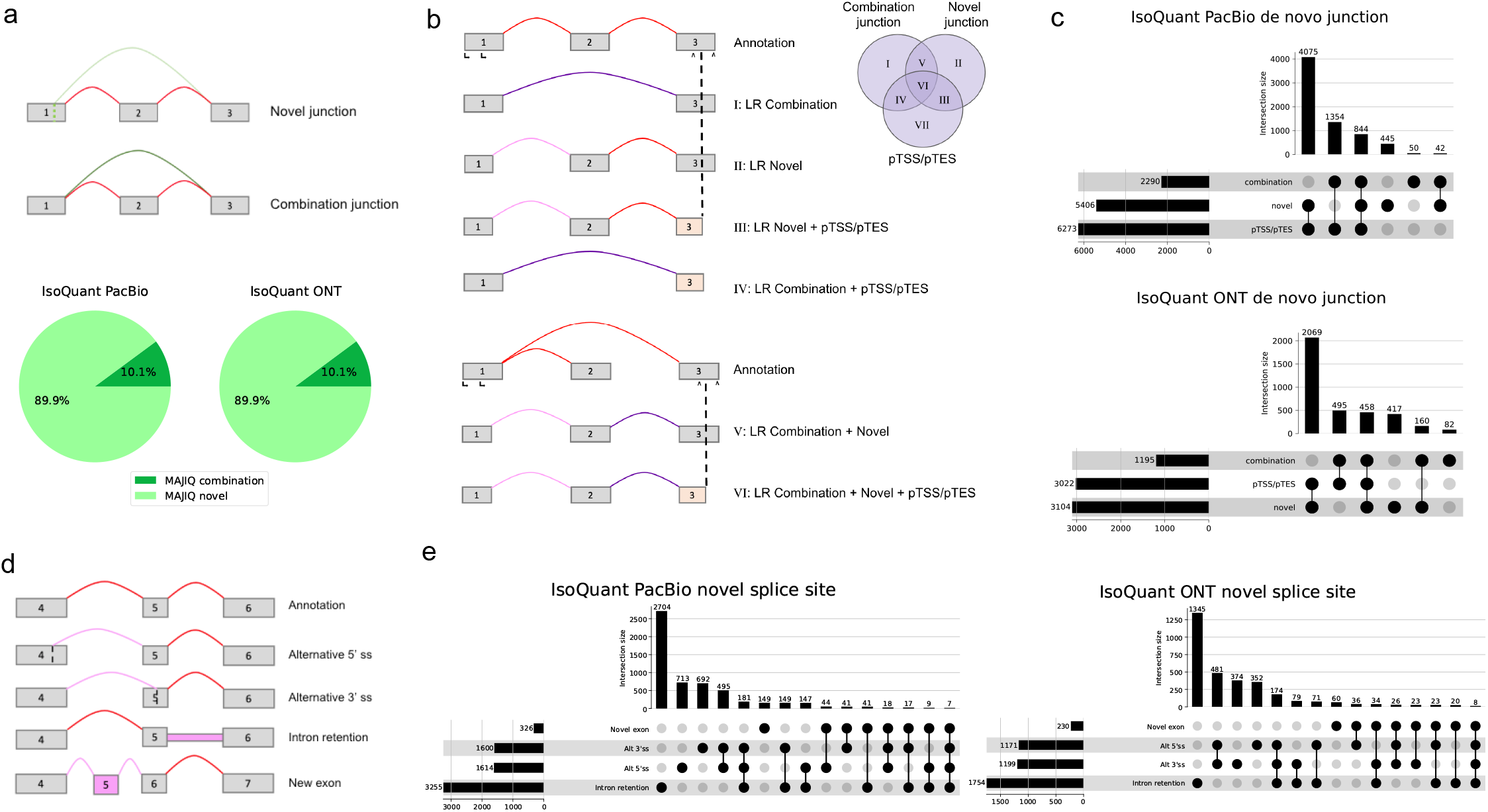
Analysis of *de novo* elements. **(A)** Short reads *de novo* splice junctions reported by MAJIQ (green junctions in the splice graphs) can be classified as those involving novel splice sites (light green) or a novel combination of known splice sites (dark green). Red corresponds to annoated junctions. The pie chart shows that compared to long reads processed with Iso-Quant, *∼* 90% of MAJIQ *de novo* splice junctions involve novel splice sites. **(B)** Representative cartoon examples for six different categories of long reads *de novo* transcript variations. Novel combination junction (dark purple), junctions involving novel splice sites (light purple), and junctions supported by annotation (red) are are the same as in (a). putative start or end (pTSS/pTES) (light yellow), or partial exons, represent cases when the transcript start site or transcript end sites do not match those in the annotation, which happens in the first or last exon of the transcript. We note that cases involving *only* pTSS/pTES (class VII in the Ven Diagram) are not included in downstream analysis as those are not handled by MAJIQ or similar short reads based splicing algorithms so can not be directly compared. **(C)** Breakdown of all cases involving *de novo* junctions reported by IsoQuant using either PacBio (top) or ONT (bottom) long reads. Notably, almost all of those cases also include pTSS/pTES. **(D)** Representative cartoon examples for the types of novel splice variations (pink) that a novel splice variant in long reads can introduce compared to the annotation (top graph, red junctions). **(E)** Breakdown of long reads novel splice junctions (light purple in (B)) into the four different categories shown in (D) when using IsoQuant to analyze PacBio (left) and ONT (right) matched reads.

In long reads *de novo* junctions, we observe another type of transcript change, termed putative transcript start or end site (pTSS/pTES). pTSS/pTES occur when only a partial exon is being output at the edge of the transcript processed by the long read algorithm (*e.g.*, IsoQuant). We term those pTSS/pTES as these do not match annotated transcript start site (TSS) or transcript end site (TES) and can thus be either a technical artifact or a bonafide TSS/TES missing in the annotation. All possible combinations of novel splice junctions with pTSS/pTES are shown in Fig 3b, along with their matching Roman numerals in the associated Venn diagram. Notably, short reads based splicing algorithms such as MAJIQ are generally unable to call transcript start/end sites so comparison of such cases is not feasible. For this reason, we do not analyze here long reads based transcripts that only involve pTSS/pTES (category VII in Fig 3b). Nonetheless, we see that when analyzing long reads transcripts with novel splice junctions, most of those involve novel splice sites and almost all of them also include pTSS/pTES (Fig 3c).

We examined long reads novel splice sites and further categorized those into splice sites that involve alternative 5’/3’ splice sites, intron retention (IR), or new exon (Fig 3d). We find most of the novel splice sites reported by long reads involve intron retentions, with PacBio and ONT reads yielding 3,255 and 1,754 such cases respectively (Fig 3e). Novel alternative 5’/3’ splice sites or a mix of those were significantly less common, and novel exons were quite rare, only about 10% of the number of IR cases. These results, shown in Fig 3c, 3e are the average values of three replicates of PacBio and ONT shown in Fig 2b. Results from different algorithms with different datasets have similar trends except for Bambu (Supplementary Fig 3, 5). Bambu employs a precision-focused threshold called novel discovery rate (NDR) to approximate the proportion of novel candidates relative to the known transcripts found. Here, NDR of 0.1, the default value, was chosen, which means 10% of all transcripts passing the threshold are novel candidates. Increasing the rate of NDR would increase the intersection sizes in the Upset plots, but it will increase false positive cases as well^5^. We chose 0.1 because this indicates at least 90% of transcripts with a similar score are annotated, providing an intuitive precision estimation. For completion, we also include Pie chart and upset plot results for the original data where long reads have significantly more coverage than short reads data, exhibiting similar trends as well (Supplementary Fig 4, 6).

### Coverage and 3’ bias lead to differences between short and long reads transcriptome views

Splice site disagreement between PacBio and ONT was previously shown to frequently represent small shifts in splice site calls, possibly due to technical artifacts^18^. Thus, we wanted to check if a similar trend can be observed for the significant differences in splice sites detected by short and long reads. For this, we defined ‘fuzzy matching’ such that splice sites found by long read algorithms are matched with splice sites reported by MAJIQ from STAR short read alignment if those are within a certain window size apart. Then, if a splice site found by long read algorithms still cannot be matched to short reads within the given window sizes it is compared to any additional annotated splice sites. Using this ‘fuzzy matching’ approach we increased the window size from 3 bp to 8 bp in both 5’/3’ splice sites for both long reads technologies and documented the resulting changes in splice sites reported by each technology and the annotation. The results of this analysis for IsoQuant are summarized in Fig 4a where the colors of the bars match those in Fig 2b. As expected, the number of splice sites matching the annotation and captured by both IsoQuant, and MAJIQ (‘All’, blue bars) increase as the window size grows from 3 to 8, while the most significant decrease is in splice sites that are in the annotation but reported only by IsoQuant (olive bars) or sites reported only by MAJIQ (burgundy bars). These cases may represent an error in short reads or situations where short reads accurately call splice sites slightly different from the annotation (*e.g.*, NAGNAG). Regardless, the overall effect of applying the ‘fuzziness’ matching was minute. For example, only 168 splice sites changed in the window size of 8 bp among the 55,000 cases in All of IsoQuant PacBio (Fig 2b), representing 0.3% changes in matching. Similar patterns are observed in different algorithms, but FLAIR and ESPRESSO *de novo* cases decrease more compared to IsoQuant and Bambu, pointing to possible differences in precision between the algorithms in *de novo* event detection (Supplementary Fig 7).

**Fig. 4:**
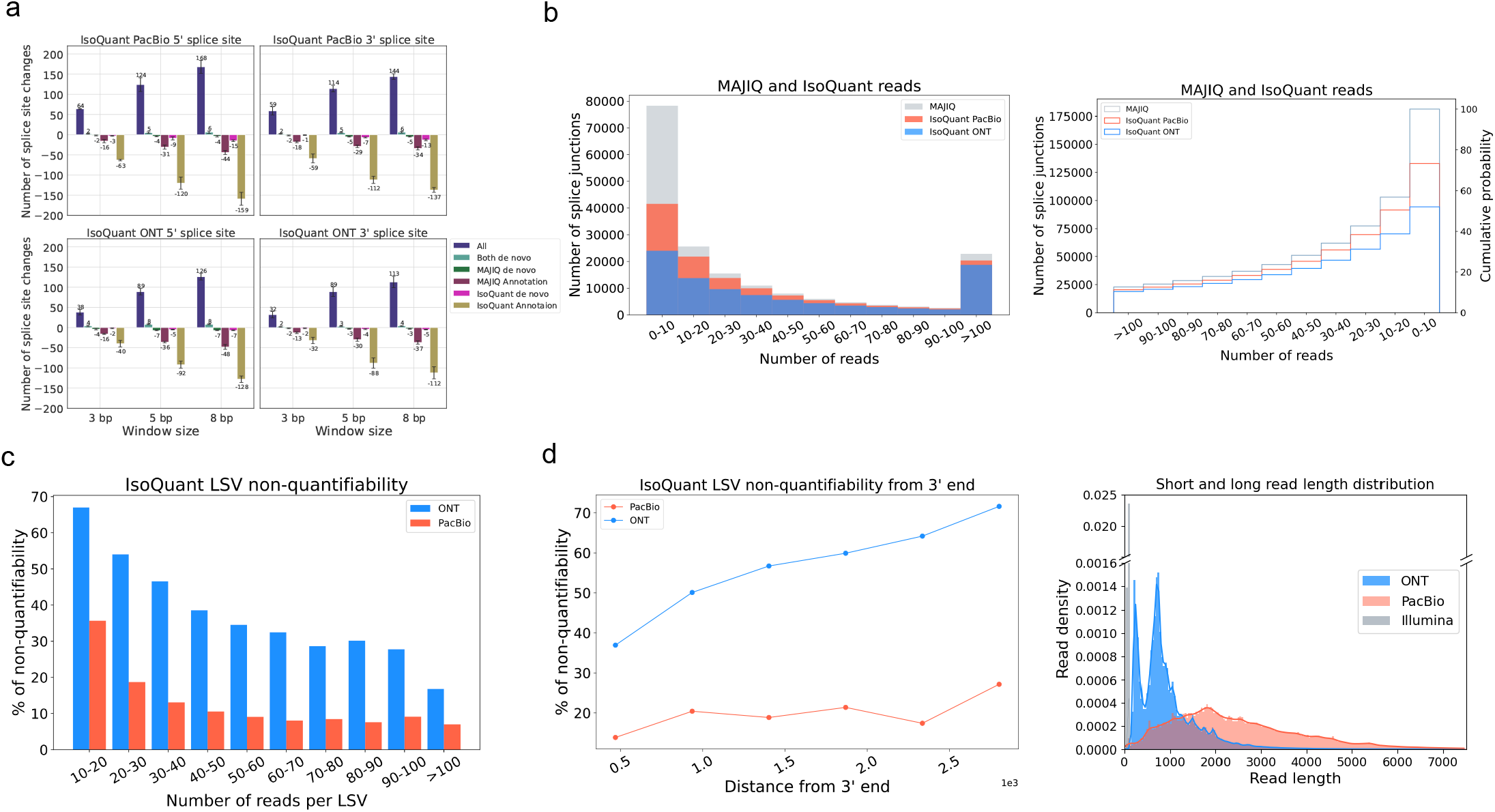
Analysis of sources of discrepancy between short and long reads based transcriptome variations. **(A)** Changes in the number of splice sites associated with each category (color) in Fig 2 as a function of ‘fuzzy’ matching window size. Here the window size (x-axis) represents the distance between long reads based splice sites and those reported by MAJIQ or the annotation at which they are still considered to match. As the window size increases the number of splice junctions in the categories All (blue) or Both *de novo* (light green) increases, while the categories for junctions detected only by long reads (magenta) or long reads and annotation (olive) drop. However, the total number of splice junctions switching their categories remains small when accounting for these splice site location discrepancies. Analysis here was performed with IsoQuant using PacBio (top) and ONT (bottom) matched reads. **(B)** The number of MAJIQ’s splice junctions (grey) identified by IsoQuant using PacBio (tomato) or ONT (blue) as a function of the number of short reads covering the junctions. The histogram on the left and CDF on the right show the number of splice junctions (y-axis) as a function of read number (x-axis). **(C)** Bar plots showing the fraction of LSV reported by MAJIQ’s short reads analysis, which were ‘non-quantifiable’ by IsoQuant using PacBio (orange) and ONT (light blue) matched long reads data. Here a ‘quantifiable’ LSV require at least 10 reads covering its respective junctions. Of note, a substantial fraction of LSV remain unjustifiable by long reads even for those with extremely high short read coverage (*>*100 reads). **(D)** Same plot as in (C) for the fraction of non-quantifiable LSV by long reads data, but here as a function of distance from transcript 3’ end. When LSV involved transcripts with multiple 3’ ends, the shortest distance was used as a conservative estimate. **(E)** The length distribution of short and long reads in the LRGASP dataset used in all the above sub-figures. Note that (b)-(d) are averaged across the three LRGASP data replicates.

To assess the relation between splice site detection and read coverage we plotted the fraction of splice sites detected by IsoQuant as a function of the number of short reads covering the junction reported by MAJIQ. For both PacBio and ONT detection was significantly worse for lowly covered splice sites with up to 10 short reads (Fig 4b). Still, even for splice sites with high short read coverage (*>* 100 reads) PacBio and ONT missed 11% and 18% of the splice sites respectively with IsoQuant. Overall, the cumulative distribution function plot shows IsoQuant PacBio detects 74% and ONT detects 52% of MAJIQ’s total amount of junctions (Fig 4b right). Among the four long read algorithms we tested, IsoQuant recovers the most junctions and Bambu recovers the least (Supplementary Fig 8a). In summary, much of the difference between short and long reads can be explained by coverage, but a significant fraction of highly covered junctions are still not detected by long reads.

The combination of several splice junctions form AS ‘events’, which serve as the base for detecting transcriptome variations with short reads technology. We thus asked how many of such AS events, or LSV in MAJIQ’s formulation, can be quantified by matched long reads. For an LSV to be quantified we require at least 10 reads to span the LSV’s junctions, which is the default LSV quantifiable filter in MAJIQ. Fig 4c shows that for IsoQuant PacBio is unable to quantify 36% of the LSV quantified by MAJIQ when having 10 to 20 short reads, while ONT cannot quantify over 65% of the LSV in the same bin. Although the non-quantifiability decreases as the number of reads per LSV increase, even for LSV with over 100 short reads PacBio is unable to quantify 8% of the LSV, a fraction similar to the one observed for splice junctions detection above. Strikingly, the fraction of non-quantifiable LSV by ONT is higher, at 18% of LSV with more than 100 reads. These observations were consistent across all four long read algorithms (Supplementary Fig 9a).

The results described above led us to hypothesize that an important contributing factor for the observed gaps between long and short reads based splicing variations is the inherent 3’ to 5’ bias of long reads technologies. PolyA selected long reads naturally begin from the 3’ end. Their length distribution is such that only 5.4% of ONT and 51.6% of PacBio reads in the LRGASP dataset shown in Fig 4d (right panel) actually span 3000bp or more, which is roughly the median length of human transcripts^19^. Furthermore, as noted above, long reads report more novel IR. This means that any splice junctions downstream of those IR events are captured further away from the polyA tail, hence less likely to be detected. To assess the effect of the 3’ bias in long compared to short reads we repeated the analysis of Fig 4c but with LSV binned by their distance from the long reads 3’ end. Fig 4d (left panel) shows that with ONT IsoQuant can not quantify about 78% of LSV when these are more than 2,500 bp away from the 3’ end. However, approximately 38% of LSV close to the 3’ end are also non-quantifiable. For PacBio data, which offered longer reads, the distribution of non-quantifiable LSV as a function of 3’ end distance is much more flat: 29% of the LSV were non-quantifiable by PacBio reads when those were more than 2,500 bp away and 12% when close to the 3’ end. This 3’ to 5’ bias trend was consistent in all four algorithms (Supplementary Fig 9b).

### Long reads detect many more intron retention events but fewer long introns

Previous sections investigated the differences between short and long reads in terms of splice junctions. However, long reads have a natural advantage in detecting Intron Retention (IR) since a single molecule may be sufficient to call such events. In contrast, IR are not directly detected by short reads and many commonly used short read algorithms such as LeafCutter^20^ do not detect IR, or do not allow *de novo* IR events (*e.g.*, rMATS^21^). Short read algorithms that do detect IR events rely on various filters and thresholds over reads that cross the splice junction into the intron or read coverage across the body of the intron. This makes IR detection from short reads highly dependent on those filtering criteria. In the analysis below we used MAJIQ’s default parameters, which are quite conservative for IR detection^11^.

While long reads can give direct evidence for IR events, these detected IR events still raise the question of whether these are reliably detected and biologically significant. To address this, we computed PSI values for long reads IR events reported by different algorithms (see Methods for details). We also plotted the fraction of introns detected by MAJIQ as a function of the number of long reads covering the intron reported by IsoQuant.

Figure 5a shows the counts of unique IR events reported by MAJIQ’s short reads and matching PacBio and ONT long reads by IsoQuant using the LRGASP data. Here, IsoQuant PacBio reports a staggering number of *∼*10K unique IR events, compared to about 6.3K with ONT, and 2.4K unique MAJIQ short read based IR events. Investigating these IR event sets, we find MAJIQ’s unique set to include longer introns (Fig 5b). This result is to be expected given the limitation of long reads overall length and 3’ bias discussed above. Finally, when we assess the relation between IR events detected by IsoQuant using either PacBio or ONT reads and IR events detected by MAJIQ from short reads, we find no direct relation between detection and IR PSI value or number of reads. This result agrees with previous reports regarding the limitations in IR detection from short reads. Of note, the results shown in Fig 5a-c, are the average values of three replicates in LRGASP data, and similar trends were observed when we used different long reads algorithms (Supplementary Fig 10, 11).

**Fig. 5:**
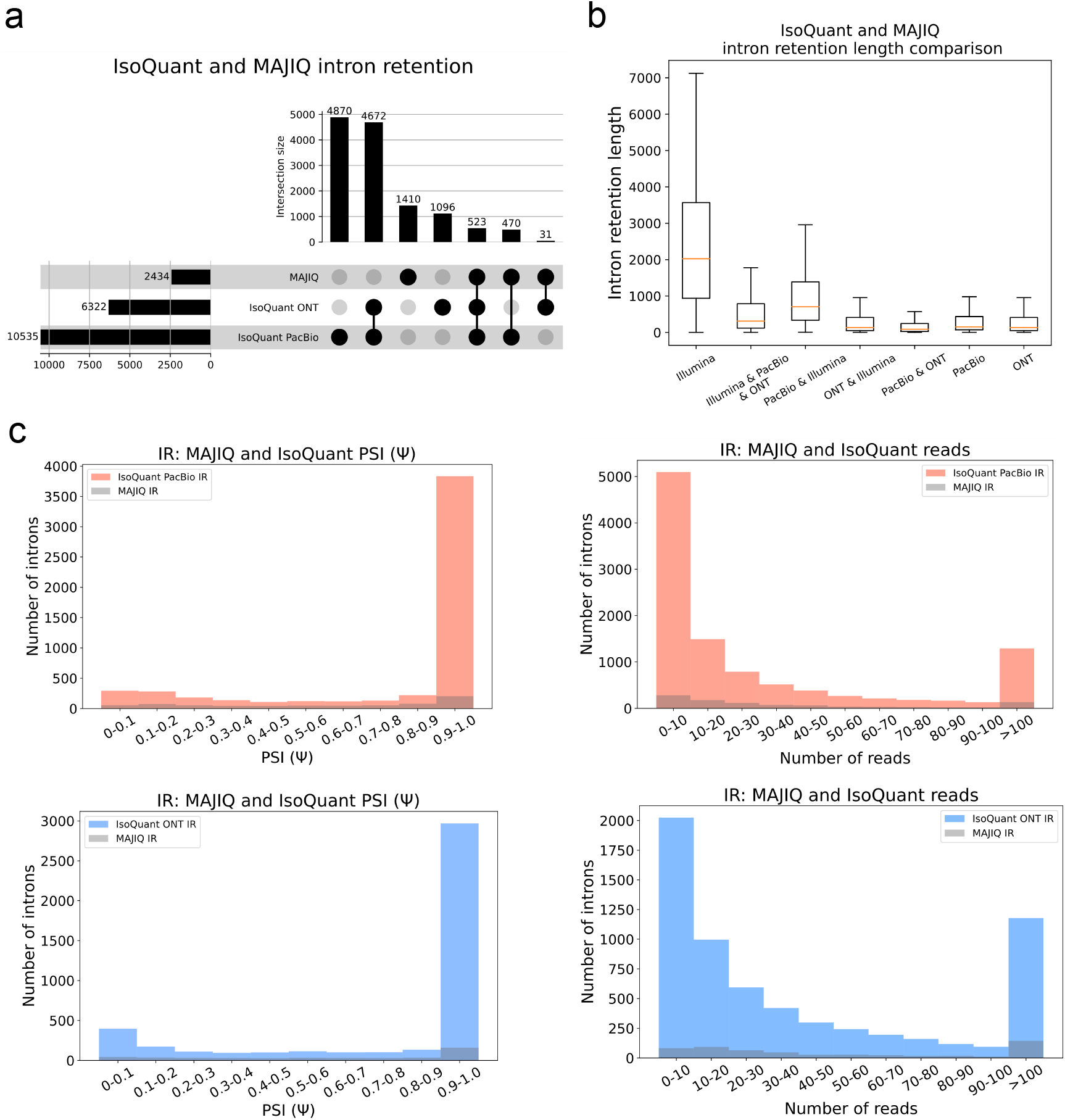
Comparison of intron rention (IR) events. **(A)** Upset plot showing overlap and total IR events reported by MAJIQ from short reads and IsoQuant using PacBio or ONT matched long reads (LRGASP dataset). **(B)** Boxplots showing IR length distribution across seven categories in Fig 5a. Each boxplot represents the IR length (y-axis) in each category (x-axis). The median is denoted by the yellow line, the upper and lower quartiles are denoted by the box, and the whiskers show points that lie within 1.5 IQRs of the lower and upper quartiles. The number of events in each category corresponds to those in (A). **(C)** Introns reported by IsoQuant using PacBio (tomato) or ONT (blue) and how many of those were also identified by MAJIQ (grey) as a function of the PSI values (left) or the number of reads covering the intron (right). For PSI, only IR events with at least 10 reads were considered and if an intron appears multiple times, the lowest PSI value is chosen for it.

### A Unified Visualization of Short and Long Read-Seq with VOILA

The comparative analysis of transcriptome variations clearly pointed to the complementarity between short and long reads RNA sequencing. In order to facilitate unified visualization and downstream analysis of short and long reads we developed the VOILA V3 package. VOILA V3 is able to combine MAJIQ’s short reads splicing analysis with GTF output files from any long read algorithm. An illustrative example is shown in Fig 6, where we ran MAJIQ with short reads and IsoQuant with PacBio long reads on one of the human cell line samples in LRGASP^14^. VOILA v3 can show short reads based gene splicegraph (first row), a unified short and long reads gene splicegraph (second row), and a list of transcripts found by only long reads (row three and below). Each color for splice junction and intron retention shows what source supports it (short, long, and annotation), with colors matching those in Fig 2b. Users can filter their data by several criteria including which source the splice graph element came from, read coverage over junctions, LSV types and complexity. Distributions over PSI for both short and long reads are represented using violin plots as the dotted black boxes in the first and second rows (see “Methods”). For long reads, the read number per transcript, junction/IR, and exon are displayed. Also, the visualization displays TSS/TES of each transcript. For a unified splicegraph visualization, junction/IR read numbers for both short and long reads are stated, and TSS/TES are not represented.

**Fig. 6:**
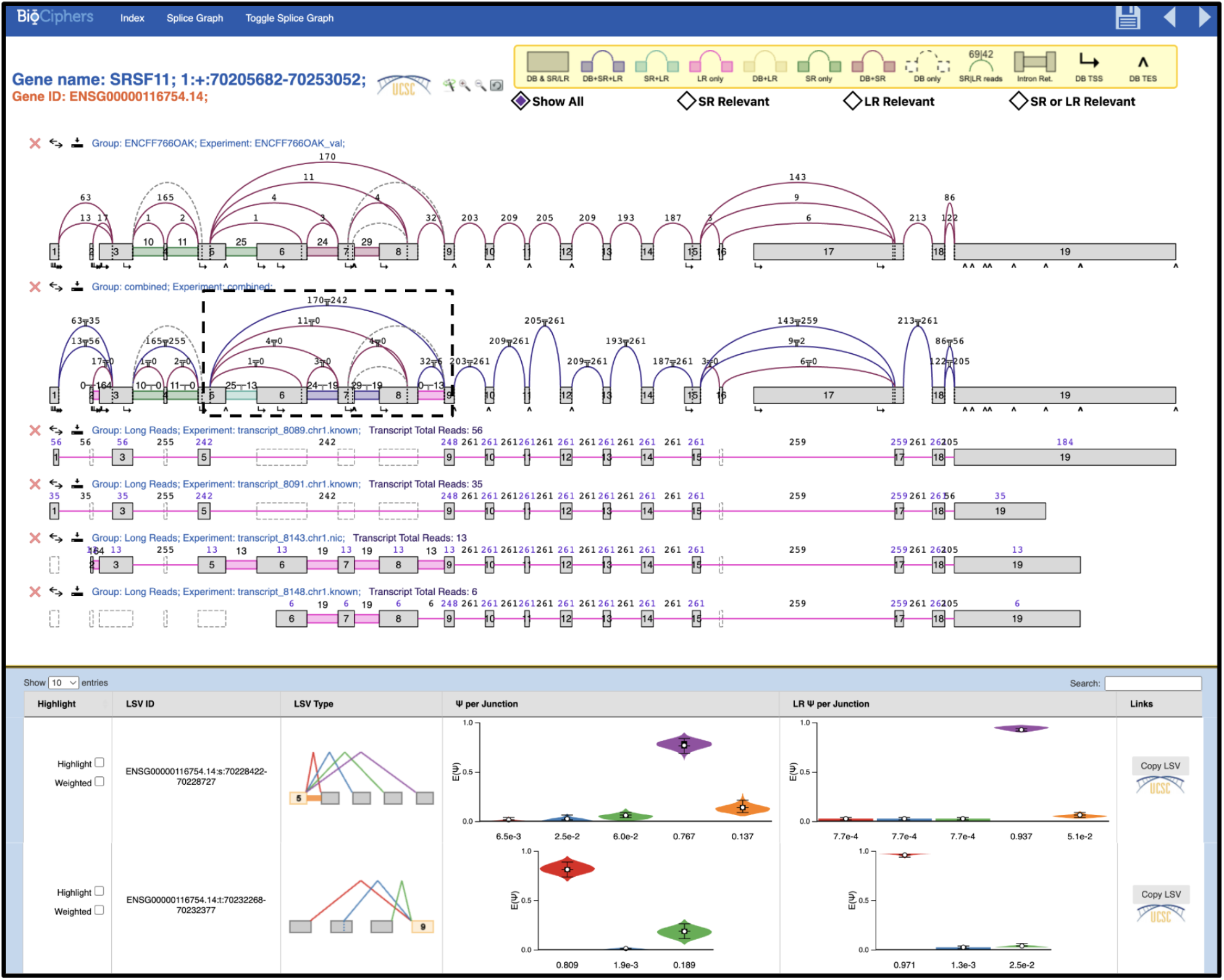
MAJIQ-L integrative analysis of splicing variations using short and long reads. Snapshot of the VOILA v3 interactive visualization of MAJIQ-L output for the SRSF11 splice factor using the LRGASP data, including short reads processed by MAJIQ and PacBio reads processed by IsoQuant. The top portion shows gene information and filtering criteria between short and long reads as well as the short reads splicegraph, a unified splicegraph, and a list of transcripts reported by IsoQuant for SRSF11. Read numbers for each transcript and the marginal count for each specific elements (junctions, introns, and exons) are included. In the unified splice graph view read count for junctions and introns are shown for both short (left) and long (right) reads, separated by a *T* sign. Note that pTSS/pTES are only shown in transcripts found by long reads. The bottom portion shows distributions of E(Ψ) values of LSV that both short and long reads find, displayed as a violin plot for the exon 5 source LSV and exon 9 target LSV in the black dotted box. The source of the individual junction can be highlighted by hovering the cursor over the junction and multiple filters can be applied interactively.

To demonstrate the usage of VOILA v3, we show the splicing analysis for the splicing factor SRSF11. The black dotted box of the SRSF11 gene in the unified splicgraph (second row) highlights alternative exons that introduce ultra-conserved premature termination codon (PTC) that induce nonsense mediated decay (NMD). Such PTC introducing exons are known regulatory feature controlling many RBPs, especially those in the serine/arginine protein family^22;23^. Both short and long reads support the identification of this important regulatory mechanism, but some differences can be observed. In general, short reads detect more diverse splicing patterns with more splice junctions coming out of exon 5, resulting in more dispersed violin plots. Both short and long reads detect multiple unannotated IR events in this region, but only long reads detect 13 reads spanning the last IR event between exon 8 and 9. These results are inline with our more general, transcriptome wide, analsyis as long reads are more likely to have a bias towards short isoforms (skipping the exons) yet can more easily detect intron retention events. Regardless of these differences, it is important to note the complex splicing patterns that emerge from this analysis. Specifically, in the context of NMD triggering alternative splicing events concerning serine/arginine proteins, the prevailing notion is that poison exon inclusion is the primary contributor to NMD. However, evidence from both short and long reads suggest that IR in this region may also play a role in controlling SRSF11 expression.

## Discussion

The work presented here was motivated by the rapid adaptation of long reads RNA sequencing. Our labs, as many others, found unique advantages to using long reads. Specifically, long reads allow researcher to resolve the relation between separate AS events along a gene splice graph and assess which isoforms include specific splice junctions of interest. Resolving full isoforms is of particular importance when trying to detect, for example, novel immunotherapy targets^24^. Similarly, the ability to unequivocally and with high sensitivity detect intron retention and overcome mappability limitations over repetitive regions are significant advantages of long reads technologies. However, the qualitative limitations of low coverage, higher error rates, combined with the fact that extensive short reads data already exists, led us to compare and contrast the short and long reads based transcriptome variations detection and quantification. Specifically, we formulated three questions as the base for this study: How to compare/contrast transcriptome variations detected by short/long reads, is there utility in combining those, and if so can we develop a method for such a combined analysis?

To address the first question we formulated a set of metrics by which any long reads technology or algorithm can be compared to MAJIQ’s short reads based splicing anlaysis. We showed there is a significant gap in both detection and quantification of splicing variations. Short reads based analysis of matched datasets revealed 40% to 60% more splice junctions, with PacBio detecting approximately 10% more than ONT. 11-18% junctions with high short reads coverage (*>* 100) were missed by PacBio and ONT, and 10% to 30% of splice junctions with PSI *>* 20% were missed by PacBio and ONT respectively. As for the ability to quantify local splicing variations, we found a clear 3’ to 5’ bias with 12-29% (PacBio) and 38-78% (ONT) of the LSV quantified by MAJIQ from short reads were unquantifiable by IsoQuant as a function of the distance from the polyA tail. This phenomena is not unique to IsoQuant and seem to reflect the limited length of the long reads, especially those in the ONT datasets we analyzed here.

On the other hand, IsoQuant PacBio and ONT were able to detect significantly more unique IR events, respectively 10K and 6.3K, compared to *∼*2.4K for MAJIQ short reads IR. These results point to the complementary nature of currently available short and long reads assays and we developed a software package, MAJIQ-L, to enable such a combined analysis.

We believe the significance of the work presented here stems from several factors. First, our results clearly demonstrate the benefit of a combined short and long reads analysis. Second, we are painfully aware that long reads RNA sequencing is a fast evolving technology with more algorithms and improved protocols or assays frequently announced by researchers or companies. Thus, a second significant component of this work is that the pipeline we developed here can be used to independently assess any newly released long reads algorithm or technology either by the developers themselves or interested users. Naturally, the introduction of new results or new technology tend to suffer from a strong confirmation bias, focusing on what is new or improved. For transcriptomics, researchers may consequently conclude long reads subsumes short reads data. However, the picture we draw here is more complex. Thus, we view the ability to easily and independently compare technologies’ output using our pipeline, as well as the results we already provide, as key for genomics researchers to make informed decisions. A third significant component of this work is MAJIQ-L with the VOILA V3 visualization package to allow researchers to perform integrated long and short reads analysis.

Our analysis points to several key conclusions. First, to answer many scientific questions researchers may be best served by short and long reads combined analysis, possibly utilizing existing short reads data, and a two stage approach - initial discovery with short reads, then focused targeted sequencing with long reads of specific genes of interest. The higher costs of long reads sequencing further points to the utility of such an approach. Moreover, our results clearly show that costly deeper long reads sequencing alone may not be an effective solution. Rather, researcher who want a more complete transcriptomic view from long reads should aim to extend the length of the long reads. We note that this result is directly coupled with the library preparation protocols typically used in the application of each sequencing technology: Random primed, dUTP stranded protocol for Illumina, vs a template switching one which is the standard long reads cDNA protocol. While direct RNA sequencing protocols are available and have been shown to avoid RT artifact such as ‘falsitrons’^25^, the length of the consequent reads remain similar. Thus, improving the length distribution or coverage across transcripts body remain an important direction for future technology improvement.

There are several limitations and possible extensions to this work. First, MAJIQ-L with VOILA V3 allows for an integrated splicing analysis of short and long reads, but does not include a unified probabilistic model for those. Such a unified model could potentially further improve isoform level quantification. Second, there is still a clear need to resolve the many pTES/TSS sites reported by long reads as has been noted by other recent work^26^. Future extensions can also include allele specific splicing and detection of variants directly from the long reads data. We are excited to explore these directions in the future and hope the combined comparative analysis pipline and results, along with the MAJIQ-L package would be highly useful for Genomics researchers focused on transcriptome variations.

## Data and Code Availability

The matched human cell line dataset can be accessed through the LRGASP https://www.gencodegenes.org/pages/LRGASP/. The GTEx v9 heart atrial appendage is available at GTEx website https://www.gtexportal.org/home/datasets. MAJIQ-L will be part of a new version of the MAJIQ software, which will be made available at https://majiq.biociphers.org/ with a matching user support group at https://groups.google.com/g/majiq_voila?pli=1.

## Acknowledgements

We thank Danielle Gutman and Matthew Gazzara for helpful comments on the manuscript.

