## Supplemental file for "Contrasting and Combining Transcriptome Complexity Captured by Short and Long RNA Sequencing Reads"

### Supplementary Figures

a

| | MAJIQ (short read) | LR algorithm (long read) | Annotation |
|---|---|---|---|
| All | ✓ | ✓ | ✓ |
| Both de novo | ✓ | ✓ | ✗ |
| MAJIQ de novo | ✓ | ✗ | ✗ |
| MAJIQ / Annotation | ✓ | ✗ | ✓ |
| LR de novo | ✗ | ✓ | ✗ |
| LR / Annotation | ✗ | ✓ | ✓ |

b

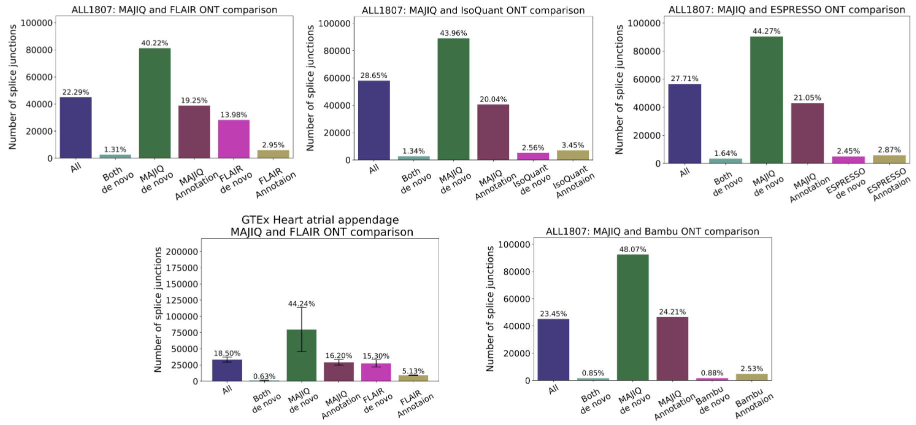


**Supplementary Fig. 1: Splice junctions comparative analysis in GTEx heart atrial appendage and PDX cell line samples. (A)** Any detected splice junction can fall into one of six categories, each represented by a color, depending which of the three sources of information (short reads, long reads, annotation) support it. **(B)** Bar charts corresponding to the aforementioned six categories. Mean and standard error bars are computed using matched datasets from three samples of human heart atrial appendage sequenced by GTEx. PDX cell line derived from a patient with a relapsed B-ALL contains only one sample. This data includes short reads processed by STAR and MAJIQ, Long reads from ONT assays, and four long read algorithms used to process the long reads data. Note that heart atrial appendage samples are only processed with FLAIR.

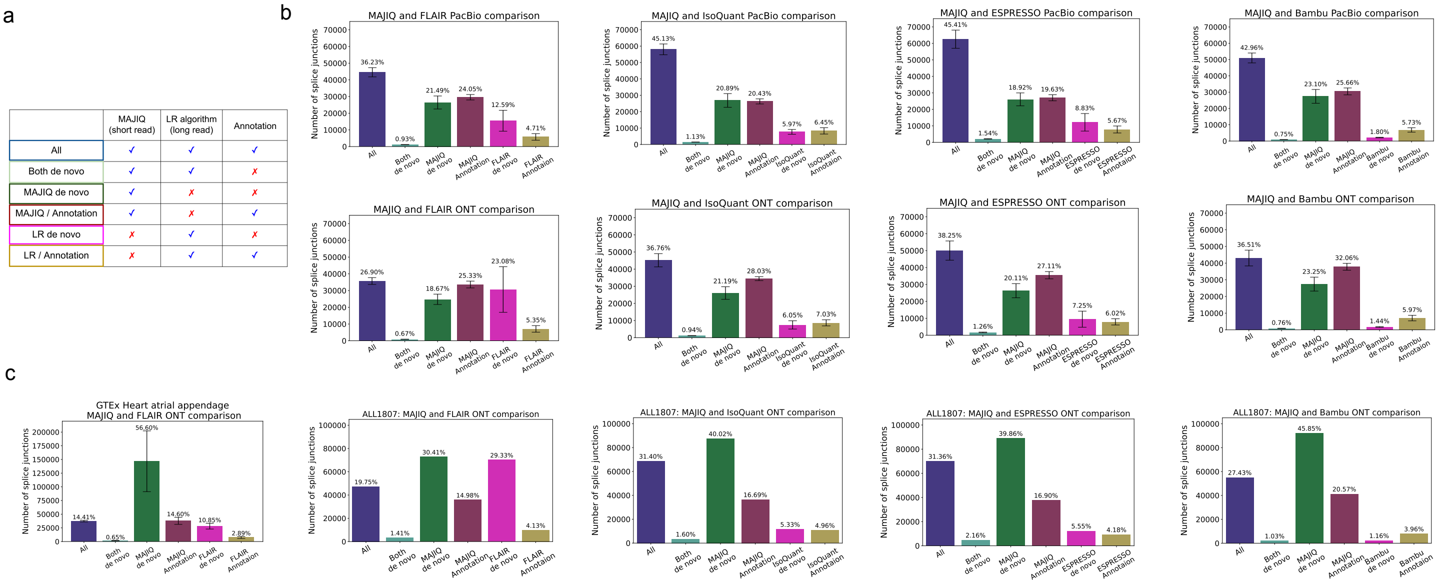


**Supplementary Fig. 2: Additional splice junctions comparative analysis when there are coverage differences between short and long reads.** **(A)** Any detected splice junction can fall into one of six categories, each represented by a color, depending which of the three sources of information (short reads, long reads, annotation) support it. **(B)** Bar charts corresponding to the aforementioned six categories. Mean and standard error bars are computed using matched datasets from three replicates of human cell line sequenced by the LRGASP14. This data includes short reads processed by STAR and MAJIQ, Long reads from PacBio and ONT assays, and four long read algorithms used to process the long reads data. Note that PacBio has 1.3-fold and ONT has 2.4-fold more coverage than Illumina in this figure. **(C)** Bar charts corresponding to the aforementioned six categories. Mean and standard error bars are computed using matched datasets from three samples of human heart atrial appendage sequenced by GTEx. PDX cell line derived from a patient with a relapsed B-ALL contains only one sample. This data includes short reads processed by STAR and MAJIQ, Long reads from ONT assays, and four long read algorithms used to process the long reads data. Heart atrial appendage samples are only processed with FLAIR. Note that ONT has 2.9-fold more coverage than Illumina in PDX cell line sample, and Illumina has 1.7-fold more coverage than ONT in heart atrial appendage samples.

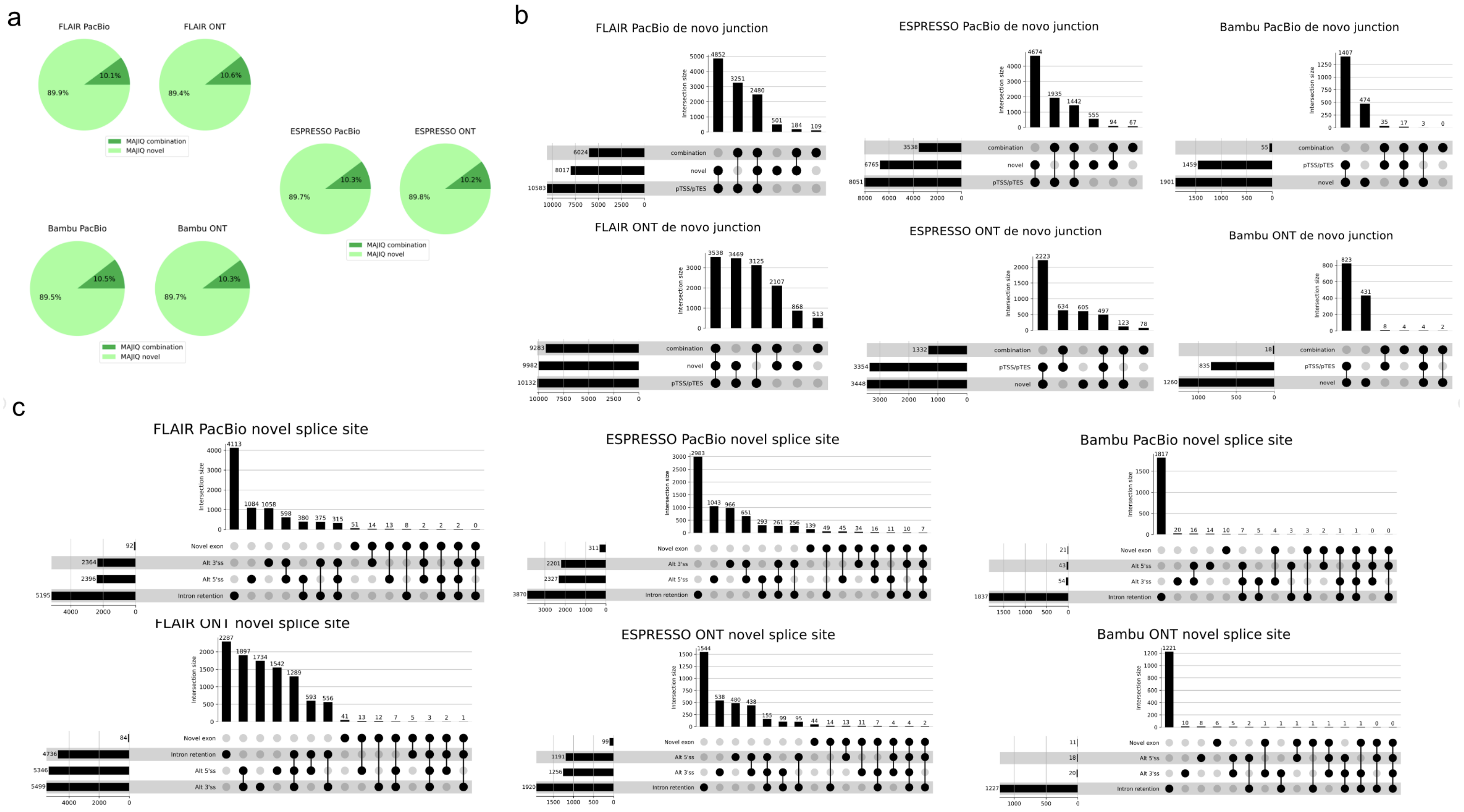


**Supplementary Fig. 3: Additional analysis of *de novo* elements in LRGASP dataset. (A)** Short reads *de novo* splice junctions reported by MAJIQ (green junctions) can be classified as those involving novel splice sites (light green) or a novel combination of known splice sites (dark green). The pie chart shows that compared to long reads processed with FLAIR, ESPRESSO, and Bambu, $\sim 90\%$ of MAJIQ *de novo* splice junctions involve novel splice sites. **(B)** Breakdown of all cases involving *de novo* junctions reported by FLAIR, ESPRESSO, and Bambu using either PacBio (top) or ONT (bottom) long reads. Notably, almost all of those cases also include pTSS/pTES. **(C)** Breakdown of long reads novel splice junctions (light purple in Fig 3-b) into the four different categories shown in Fig 3-d when using FLAIR, ESPRESSO, and Bambu to analyze PacBio (top) and ONT (bottom) matched reads.

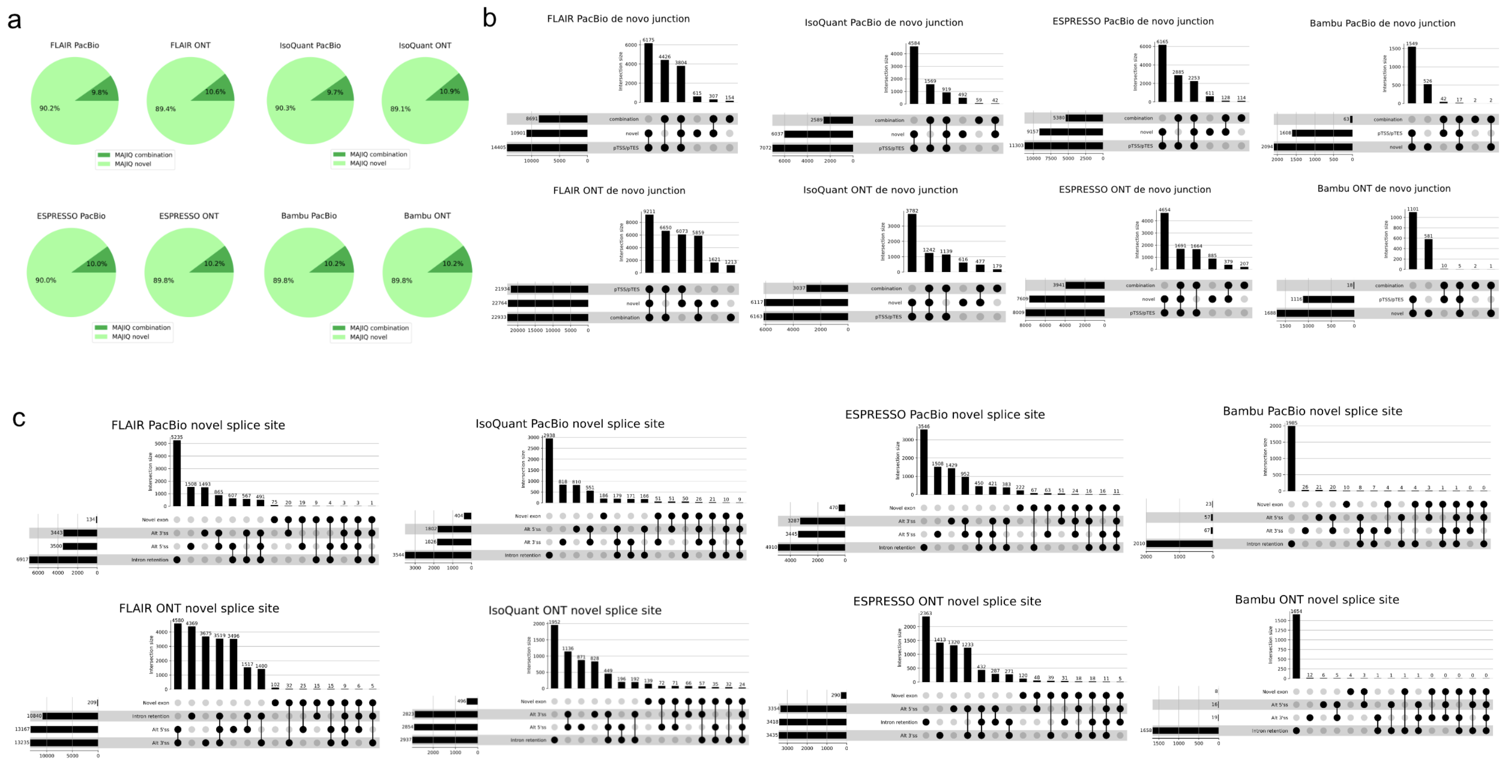


**Supplementary Fig. 4: Additional analysis of *de novo* elements in LRGASP when there are coverage differences between short and long reads.** **(A)** Short reads *de novo* splice junctions reported by MAJIQ (green junctions) can be classified as those involving novel splice sites (light green) or a novel combination of known splice sites (dark green). The pie chart shows that compared to long reads processed with IsoQuant, FLAIR, ESPRESSO, and Bambu, $\sim$ 90% of MAJIQ *de novo* splice junctions involve novel splice sites. **(B)** Breakdown of all cases involving *de novo* junctions reported by FLAIR, ESPRESSO, and Bambu using either PacBio (top) or ONT (bottom) long reads. Notably, almost all of those cases also include pTSS/pTES. **(C)** Breakdown of long reads novel splice junctions (light purple in Fig 3-b) into the four different categories shown in Fig 3-d when using FLAIR, ESPRESSO, and Bambu to analyze PacBio (top) and ONT (bottom) matched reads. Note that PacBio has 1.3-fold and ONT has 2.4-fold more coverage than Illumina in figures (a)-(c).

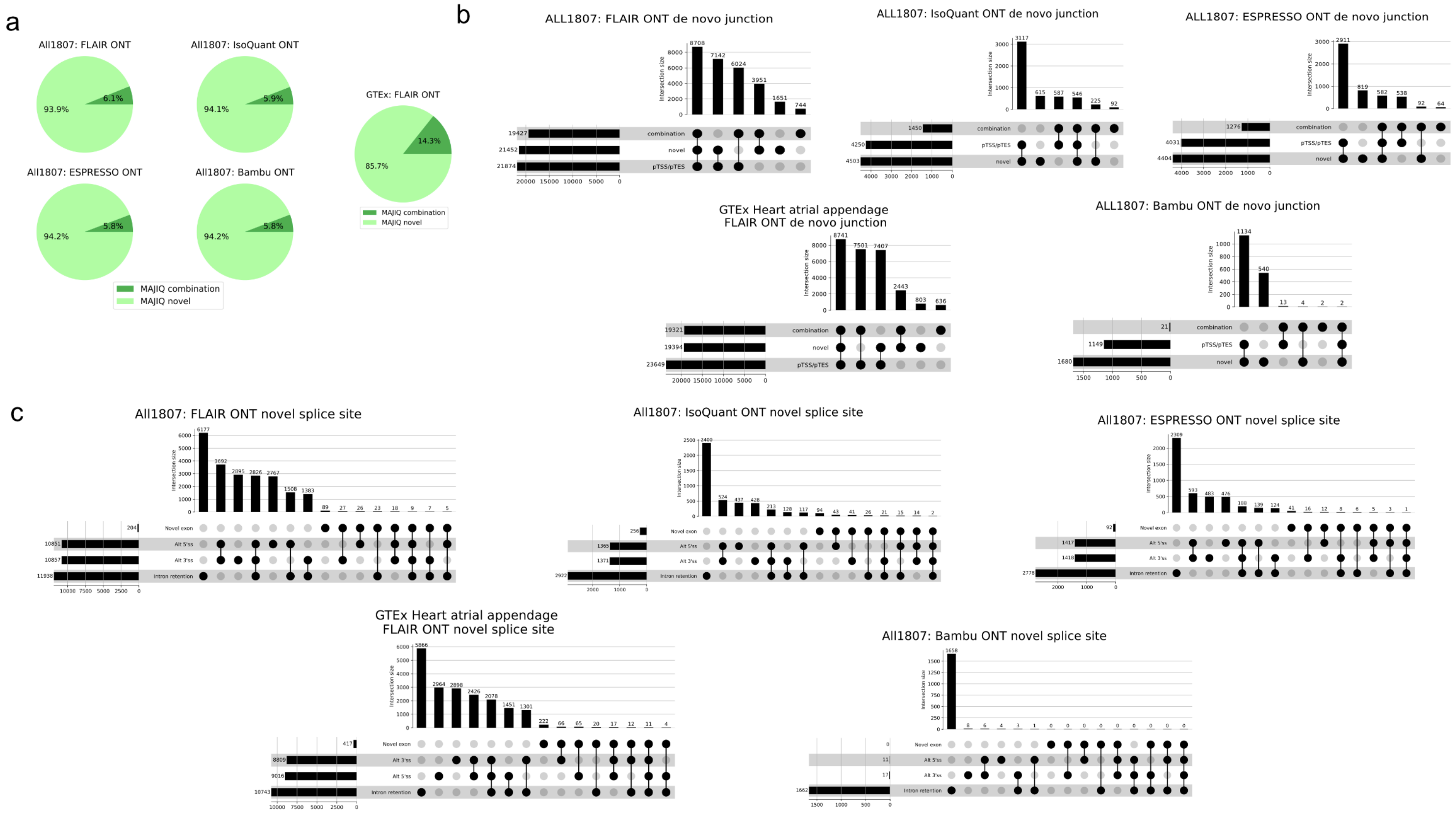


**Supplementary Fig. 5: Additional analysis of *de novo* elements in GTEx heart atrial appendage and PDX cell line samples (A)** Short reads *de novo* splice junctions reported by MAJIQ (green junctions) can be classified as those involving novel splice sites (light green) or a novel combination of known splice sites (dark green). The pie chart shows that compared to long reads processed with IsoQuant, FLAIR, ESPRESSO, and Bambu, $\sim 94\%$ of MAJIQ *de novo* splice junctions involve novel splice sites in PDX cell line sample. Compared to long reads processed with FLAIR, $\sim 85\%$ of MAJIQ *de novo* splice junctions involve novel splice sites in heart atrial appendage samples. **(B)** Breakdown of all cases involving *de novo* junctions reported by IsoQuant, FLAIR, ESPRESSO, and Bambu using ONT long reads. Notably, almost all of those cases also include pTSS/pTES. **(C)** Breakdown of long reads novel splice junctions (light purple in Fig 3-b) into the four different categories shown in Fig 3-d when using IsoQuant, FLAIR, ESPRESSO, and Bambu to analyze ONT matched reads. Note that heart atrial appendage samples are only processed with FLAIR.

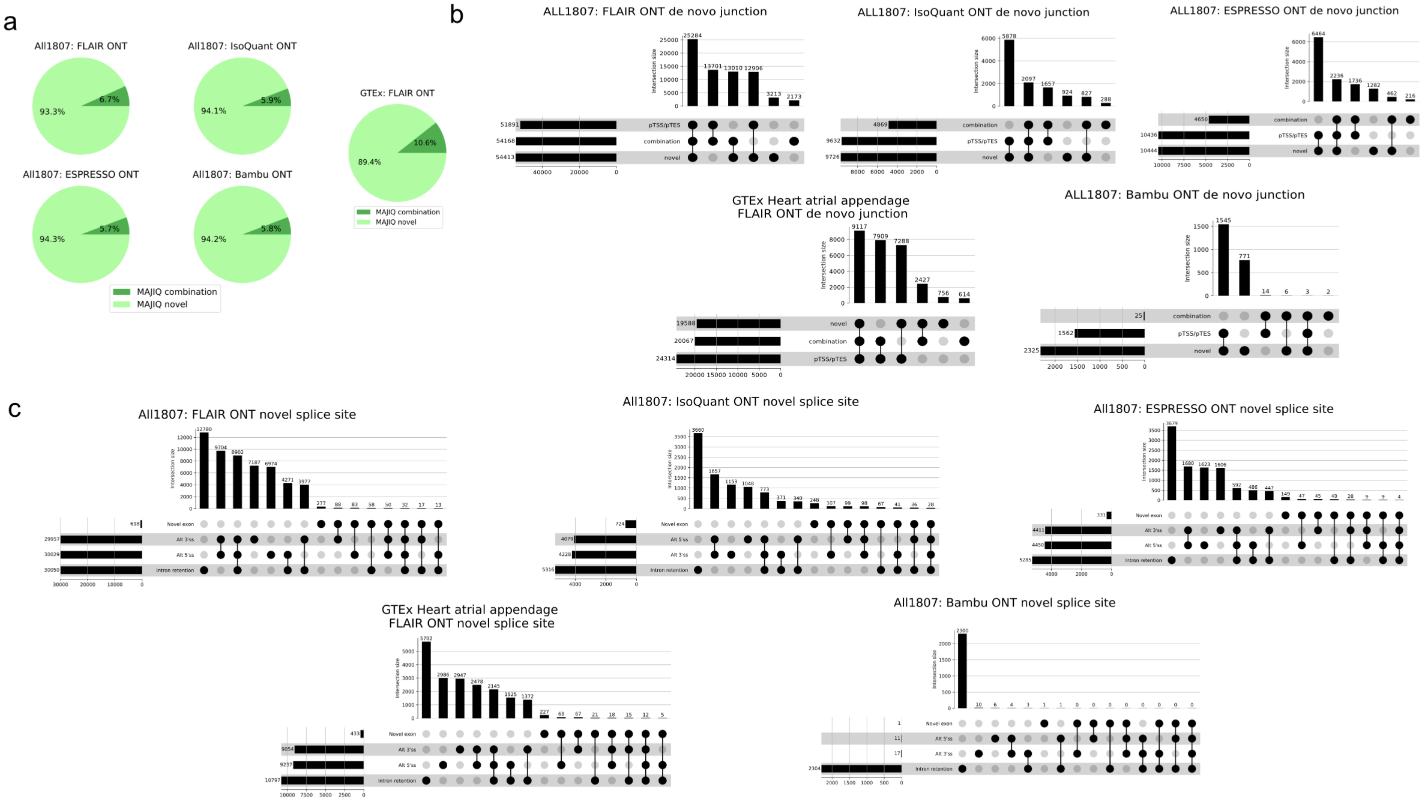


**Supplementary Fig. 6: Additional analysis of *de novo* elements in GTEx heart atrial appendage and PDX cell line samples when there are coverage differences between short and long reads. (A)** Short reads *de novo* splice junctions reported by MAJIQ (green junctions) can be classified as those involving novel splice sites (light green) or a novel combination of known splice sites (dark green). The pie chart shows that compared to long reads processed with IsoQuant, FLAIR, ESPRESSO, and Bambu, $\sim$ 94% of MAJIQ *de novo* splice junctions involve novel splice sites in PDX cell line sample. Compared to long reads processed with FLAIR, $\sim$ 90% of MAJIQ *de novo* splice junctions involve novel splice sites in heart atrial appendage samples. **(B)** Breakdown of all cases involving *de novo* junctions reported by IsoQuant, FLAIR, ESPRESSO, and Bambu using ONT long reads. Notably, almost all of those cases also include pTSS/pTES. **(C)** Breakdown of long reads novel splice junctions (light purple in Fig 3-b) into the four different categories shown in Fig 3-d when using IsoQuant, FLAIR, ESPRESSO, and Bambu to analyze ONT matched reads. Heart atrial appendage samples are only processed with FLAIR. Note that ONT has 2.9-fold more coverage than Illumina in PDX cell line sample, and Illumina has 1.7-fold more coverage than ONT in heart atrial appendage samples.

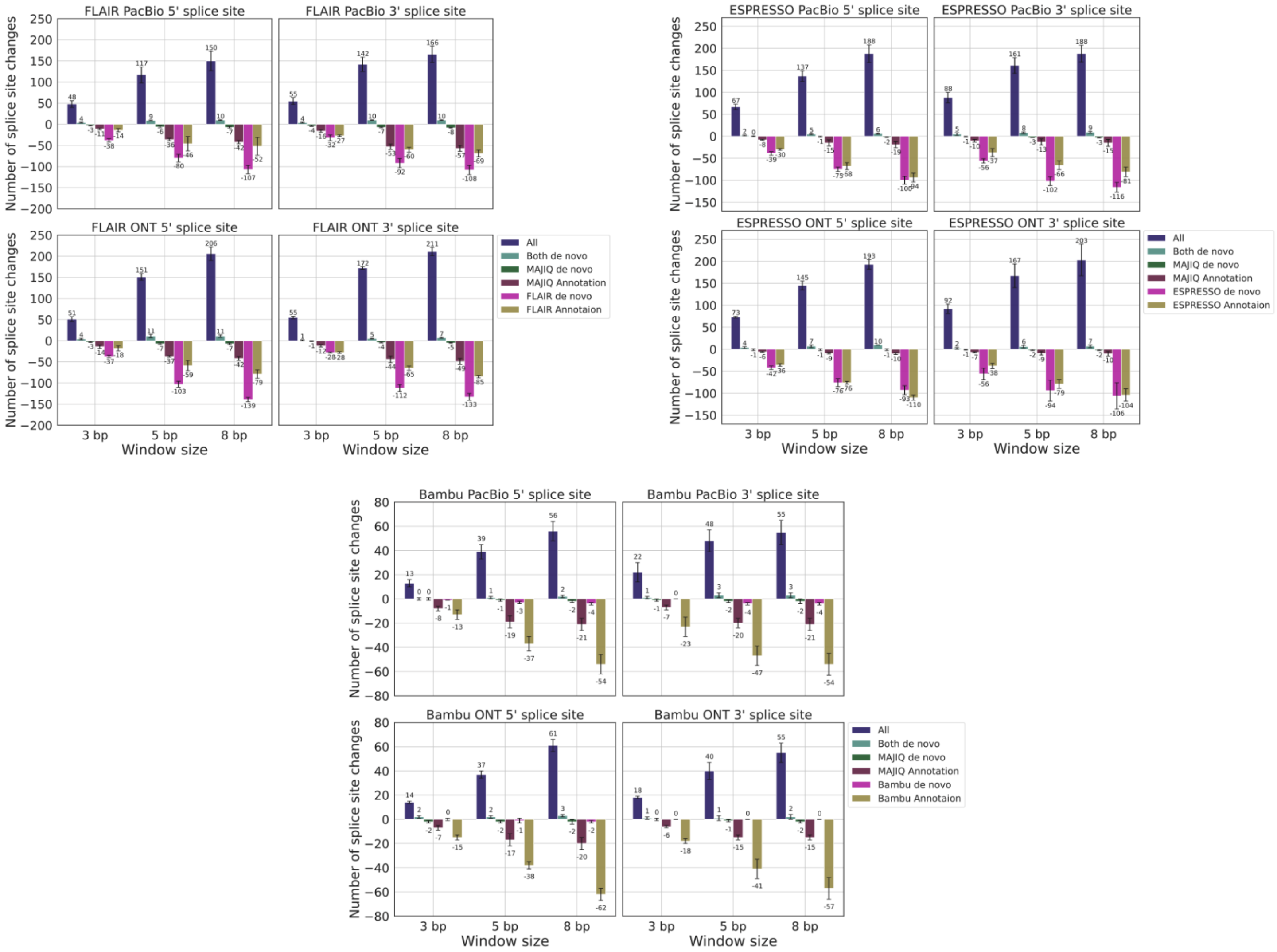


**Supplementary Fig. 7: Analysis of splice site changes in FLAIR, ESPRESSO, and Bambu.** Changes in the number of splice sites associated with each category (color) in Fig 2 as a function of 'fuzzy' matching window size. Here the window size (x-axis) represents the distance between long reads based splice sites and those reported by MAJIQ or the annotation at which they are still considered to match. As the window size increases the number of splice junctions in the categories All (blue) or Both *de novo* (light green) increases, while the categories for junctions detected only by long reads (magenta) or long reads and annotation (olive) drop. However the total number of splice junctions switching their categories remains small when accounting for these splice site location discrepancies. Analysis here was performed with FLAIR, ESPRESSO, and Bamu using PacBio and ONT matched reads.

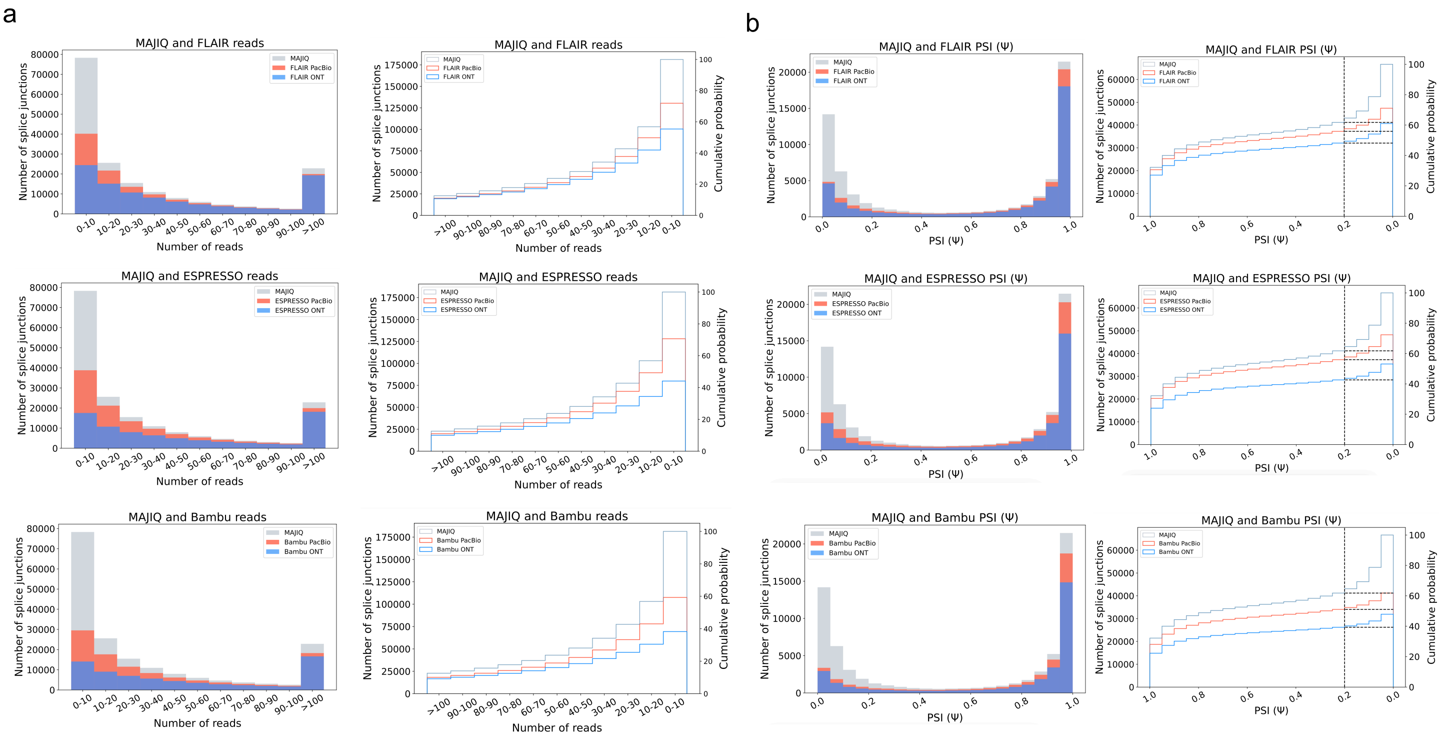


**Supplementary Fig. 8: Number of junctions identified by FLAIR, ESPRESSO, and Bambu from the junctions that MAJIQ finds.** **(A)** Bar plots showing the fraction of LSV reported by MAJIQ's short reads analysis, which were 'non-quantifiable' by FLAIR, ESPRESSO, and Bambu using PacBio (orange) and ONT (light blue) matched long reads data. Here a 'quantifiable' LSV require at least 10 reads covering its respective junctions. Of note, a substantial fraction of LSV remain unjustifiable by long reads even for those with extremely high short read coverage (>100 reads). **(B)** Taking the splice junctions reported in (B) by MAJIQ (green) and assessing the number of those also identified when using PacBio (tomato) or ONT (blue) long reads, as a function of the PSI values. Here FLAIR, ESPRESSO, Bambu were used for long reads data. Note that if a junction appears in multiple LSV, the lowest PSI values are chosen (x-axis). The graph on the right is the CDF for the histogram shown on the left. Dashed lines denote splice junctions with a PSI of 20% or more.

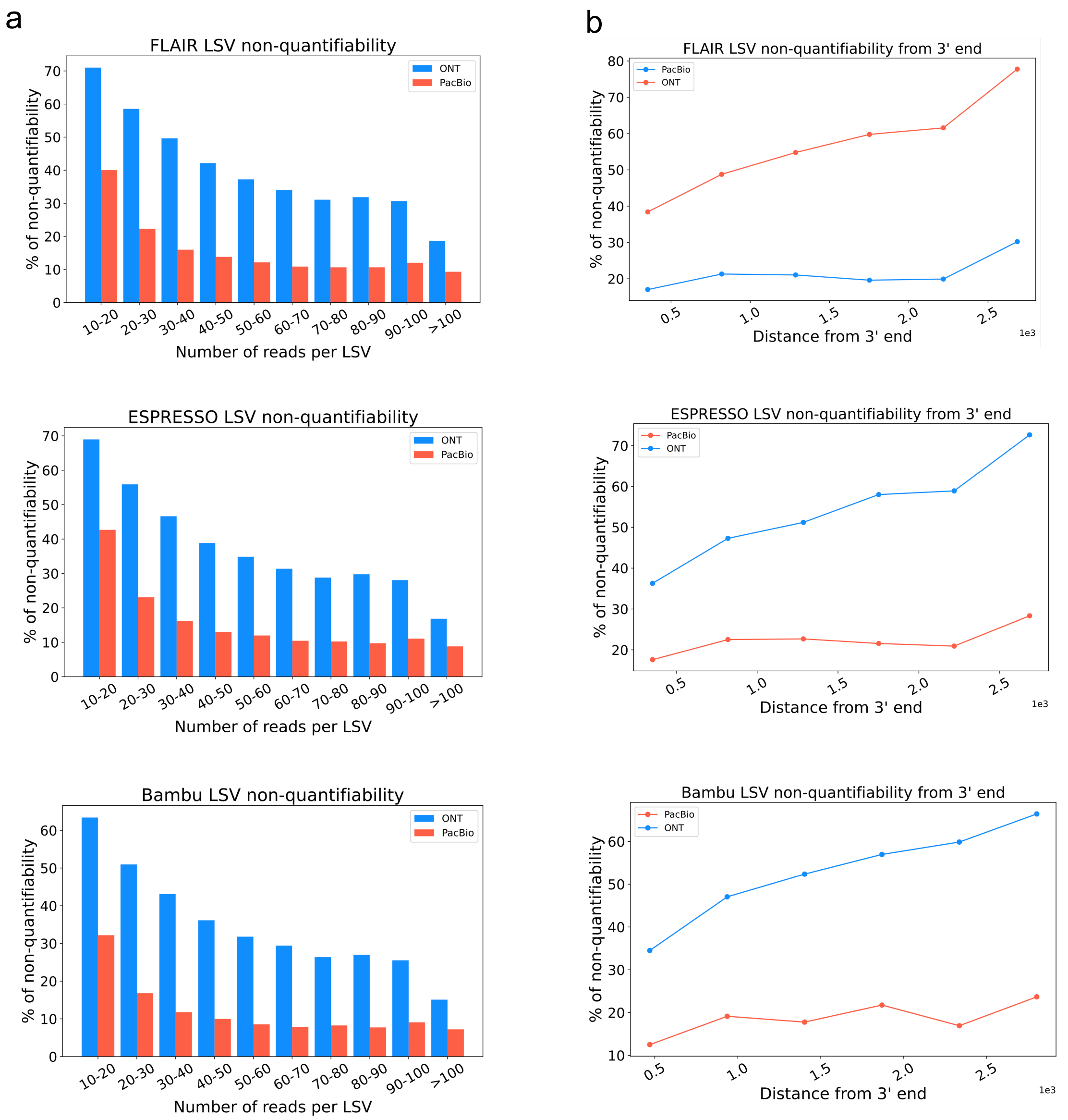


**Supplementary Fig. 9: LSV non-quantifiability and 3' to 5' bias analysis in FLAIR, ESPRESSO, and Bambu.** **(A)** Bar plots showing the fraction of LSV reported by MAJIQ's short reads analysis, which were 'non-quantifiable' by FLAIR, ESPRESSO, Bambu using PacBio (orange) and ONT (light blue) matched long reads data. Here a 'quantifiable' LSV requires at least 10 reads covering its respective junctions. Of note, a substantial fraction of LSV remain unjustifiable by long reads even for those with extremely high short read coverage (>100 reads). **(B)** Same plot as in (A) for the fraction of non-quantifiable LSV by long reads data, but here as a function of distance from transcript 3' end. When LSV involved transcripts with multiple 3' ends, the shortest distance was used as a conservative estimate.

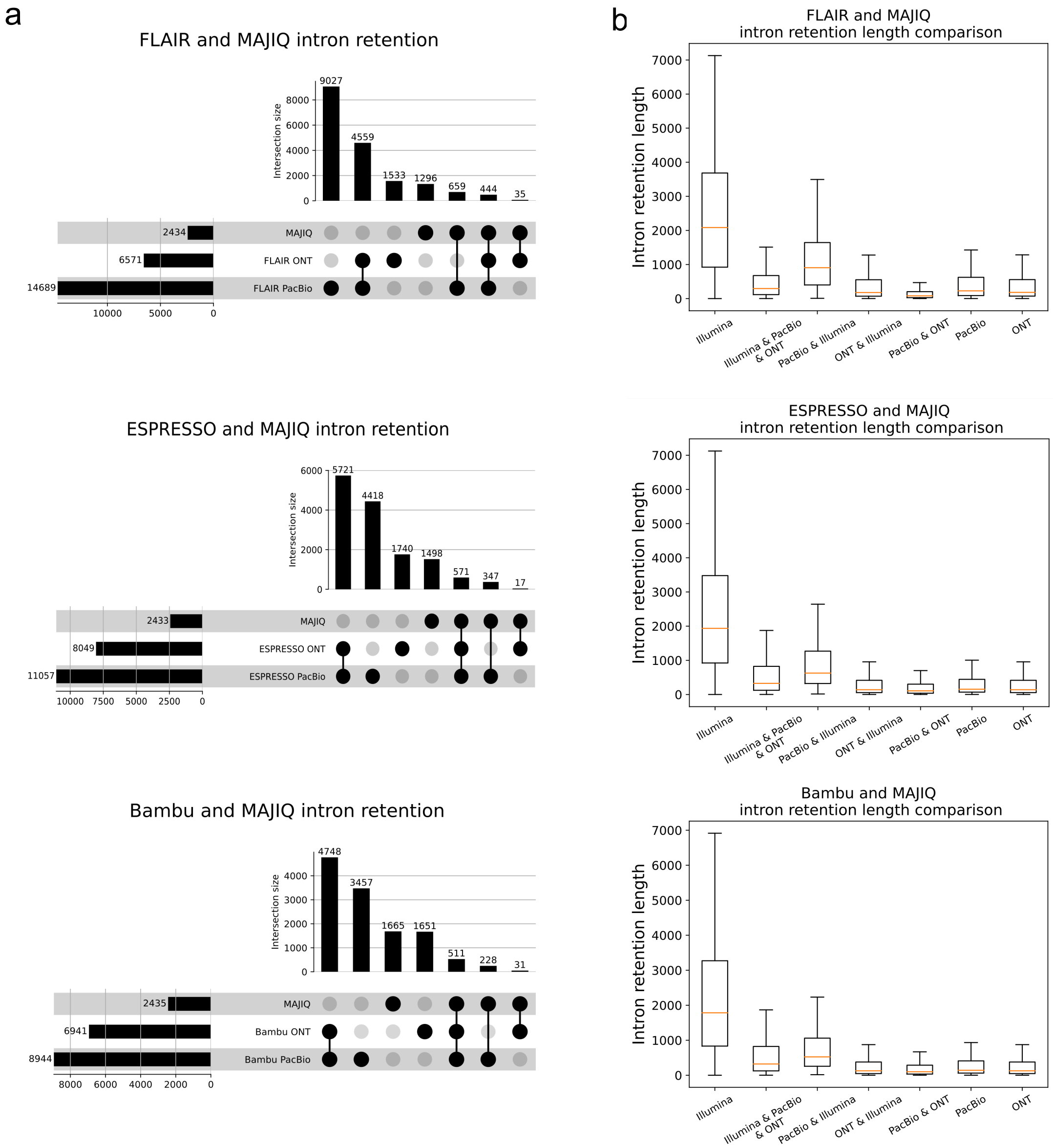


**Supplementary Fig. 10: Intron Rention (IR) events found between MAJIQ and FLAIR, ESPRESSO, Bambu. (A)** Upset plot showing overlap and total IR events reported by MAJIQ from short reads and MAJIQ and FLAIR, ESPRESSO, Bamb using PacBio or ONT matched long reads (LRGASP dataset). **(B)** Boxplots showing IR length distribution across seven categories in (A). Each boxplot represents the IR length (y-axis) in each category (x-axis). The median is denoted by the yellow line, the upper and lower quartiles are denoted by the box, and the whiskers show points that lie within 1.5 IQRs of the lower and upper quartiles. The number of events in each category corresponds to those in (A).

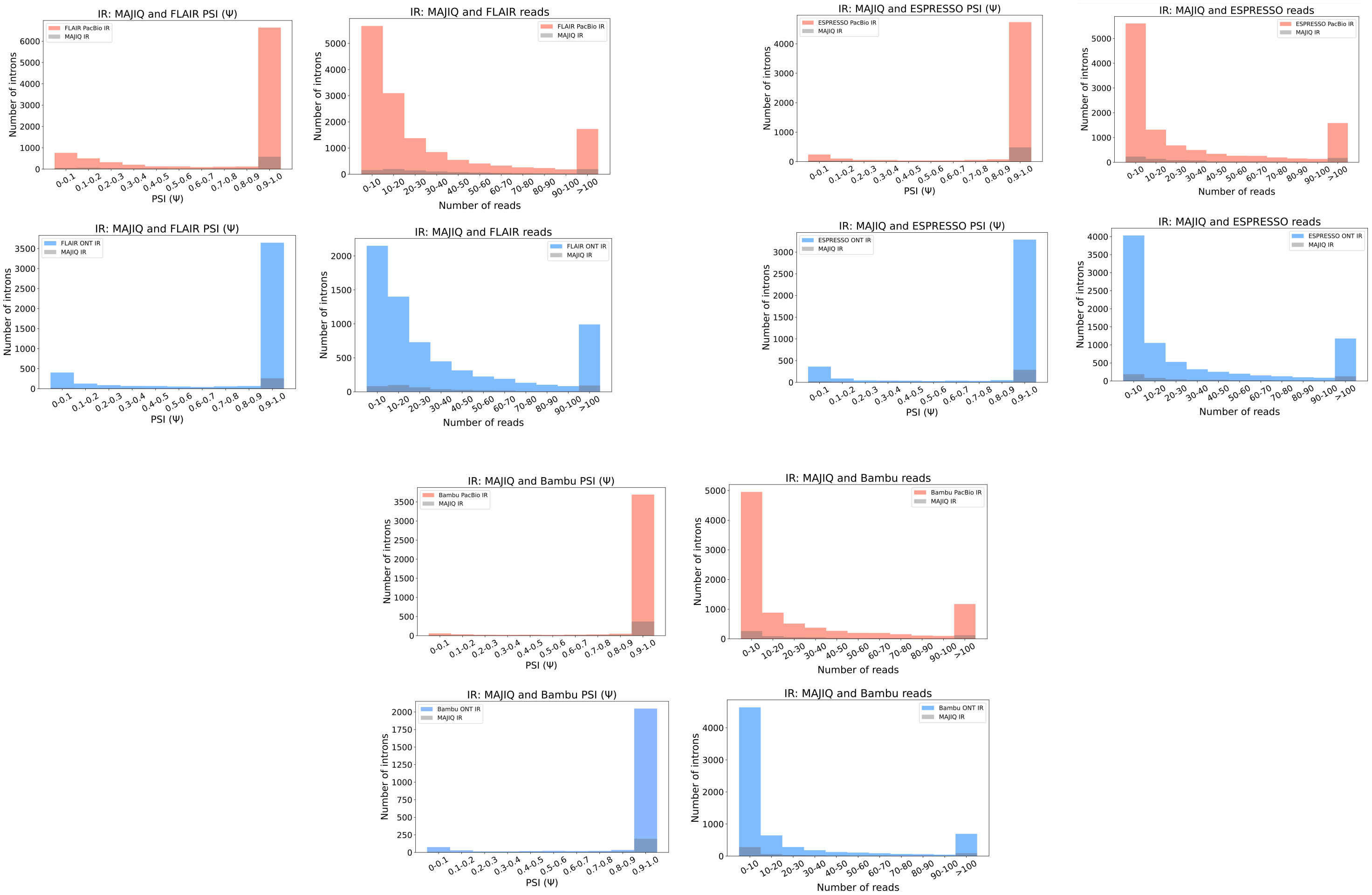


**Supplementary Fig. 11: Number of introns identified by MAJIQ from the introns that FLAIR, ESPRESSO, Bambu find.** Taking the introns by FLAIR, ESPRESSO, Bambu PacBio (tomato) or ONT (blue) and assessing the number of those also identified by MAJIQ (grey) as a function of the PSI values (left). Note that if an intron appears multiple times, the lowest PSI values are chosen. The histogram shows the number of introns (y-axis) in each PSI value (x-axis). The number of FLAIR, ESPRESSO, Bambu's introns using PacBio or ONT identified by MAJIQ as a function of the number of long reads covering the introns (right). The histogram shows the number of introns (y-axis) as a function of read number (x-axis).

### Supplementary Table

| Sample | WTC11 | | |
|---|---|---|---|
| Method | cDNA | | |
| Tech | Illumina | PacBio | ONT |
| # of replicates | 3 | 3 | 3 |
| # of reads | 137,043,475<br>(143,171,620) | 5,521,442<br>(7,424,923) | 20,888,972<br>(51,194,535) |
| Median read length | 89 | 2,209 | 610 |

**Supplementary Table 1: Coverage summary statistics of human cell line datasets in LARGASP.** For each sample, replicates were combined when reporting statistics. Note that the number inside the parentheses denotes the number of reads before subsampling. PacBio has 1.3-fold and ONT has 2.4-fold more coverage than Illumina.

| Sample | ALL1807 | |
|---|---|---|
| Method | cDNA | |
| Tech | Illumina | ONT |
| # of replicates | 1 | 1 |
| # of reads | 112,819,050 | 19,473,944 (57,523,865) |
| Median read length | 150 | 869 |

**Supplementary Table 2: Coverage summary statistics in B-ALL.** Note that the number inside the parentheses denotes the number of reads before sub-sampling. ONT has 2.9-fold more coverage than Illumina.

| Tissue | Tech | Sample | # of reads | Median read length |
|---|---|---|---|---|
| Heart atrial appendage | ONT | GTEX-1GN1W-0226-SM-7AGLJ | 5,667,159 | 630 |
| | Illumina | GTEX-1GN1W-0226-SM-7P8QY | 46,977,630 (167,645,011) | 76 |
| | ONT | GTEX-1HBPH-0226-SM-7LLUW | 8,228,317 | 718 |
| | Illumina | GTEX-1HBPH-0226-SM-9WYSM | 77,735,083 (106,764,362) | 76 |
| | ONT | GTEX-1IDJD-0226-SM-AML89 | 7,797,416 (7,955,912) | 720 |
| | Illumina | GTEX-1IDJD-0226-SM-CKZOA | 73,870,263 | 76 |

**Supplementary Table 3: Coverage summary statistics in GTEx heart atrial appendage dataset.** Note that the number inside the parentheses denotes the number of reads before sub-sampling. Illumina has 1.7-fold more coverage than ONT.

| Software | Version | Command options | PacBio | ONT |
|---|---|---|---|---|
| minimap2 | 2.24 | -ax -t 30 | splice:hq | splice |
| FLAIR | 1.7 | -t 30 | | |
| IsoQuant | 3.2.0 | --complete_genedb -t 30 | --data_type pacbio_ccs | --data_type nanopore |
| Bambu | 2.0.0 | ncore=30 | | |
| ESPRESSO | 1.3.2 | -T 30 | | |

**Supplementary Table 4: Long read tools Command line options and software versions** FLAIR, IsoQuant, Bambu, and ESPRESSO were run using the same BAM file, reference annotation, and reference genome as input.
